# Cross talk between photo-pigments and graphene electron cloud - Designing a biodiode

**DOI:** 10.1101/210609

**Authors:** Sanhita Ray, Sayantani Sen, Alakananda Das, Anirban Bose, Anirban Bhattacharya, Avishek Das, Sanatan Chattopadhyay, Hirak Patra, Shib Shankar Singha, Achintya Singha, Anjan Kr. Dasgupta

## Abstract

We report emergence of a new electrical material by growing photosynthetic biofilm on a Dirac material, graphene. The material showed new conducting as well as semiconducting properties. Frequency dependent capacitive spectra further indicated presence of electrical isosbestic points(at 0.8 and 9MHz), implying two state dieletric transitions at critical frequencies. A notable reult was a Schottky diode like behavior in the IV curve. Voltage dependent conductance with conductance peaks near the Schottky diode threshold was observed. We obtained facilitated growth of photosynthetic biofilm in presence of graphene. Lastly higher bacterial metabolism was seen in graphene incorporated biofilm. For this zero band gap Dirac material this can only be interpreted as coupling of the electron transport chain of the bacterial biofilm and the graphene electron cloud.

## Introduction

This paper explores how the zero band Dirac material^1^ called graphene is electrically modulated in presence of biofilms. The special choice of photosynthetic biofilm is made as the energy of such films is derived primarily from photon induced charge splitting. Any effect on metabolism of this bacteria would be an indicator for interactions between the Dirac material and the electron cloud associated with the electron transport chain of the bacterial species. We therefore looked into the reverse aspect, namely how beacterial metabolism is affected by presence of graphene. In most cases a biofilm inhibitory roles has been reported, but in the present case, we in fact observed a positive role as will be discussed. Graphene is a zero-band gap material. Modification of band gap in graphene has been reported by earlier groups, using standard material science approaches.^2^ We report a biofilm induced engineering of band gap in graphene. Single bacteria induced wrinkles on graphene surface and consequent emergence of Dirac point at low temperatures (10K) has been reported.^3^ However no previous workers addressed the question of biofilm induced band gap opening. This may not be totally unexpected as any close anchoring of bacterial biofilm with graphene biomass may lead to coupling between the bacterial electron transport chain and the electron cloud of graphene, a Dirac material.^1^

Biofilms are well organised microbial aggregates that are known for their resilience to adverse environmental conditions like dehydration,^4^ shear stress and chemical agents. These properties make biofilms an ideal candidate for advanced hybrid materials. Purple non-sulphur bacteria (PNSB) are capable of utilizing a wide variety of organic matter as nutrition source; at the same time it is phototropic and possesses light harvesting molecules^5^ in its membrane, that act as electron carriers. They have atypical lipid molecules in their membranes, known as hopanoids,^6^ which increase membrane rigidity. PNSB form extremely tough biofilms and along with its other properties was chosen to be a bioelectronics platform.

Lastly, we would like to bring forward some works on the application domain. Nnanofibrils and cytochromes in Geobacter biofilms^7,8^ have been investigated recently and found to possess conductive and capacitive properties, respectively. This has opened up the the possibility of using microbes in electronic devices. For microbe based electronics, conducting and/or semi-conducting nano-materials need to be incorporated into microbial consortia.

**Figure 1:**
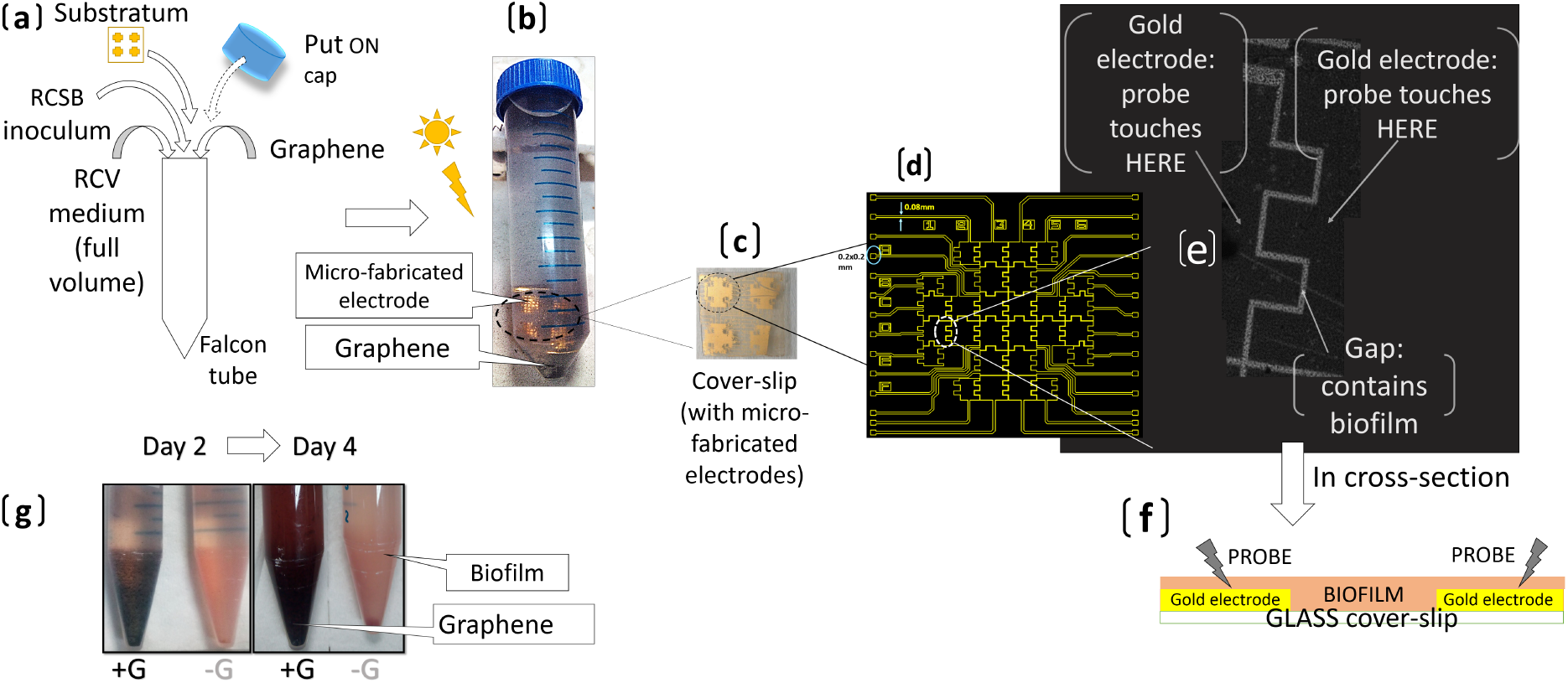
Synthesis of graphene-biofilm bio-composite. (a) Schematic of components added to bacterial growth medium and conditions for growth of bio-composite. Panel (b) shows the set-up upon inoculation. Panel (c) shows gold electrodes fabricated upon glass coverslip, 4 complexes of electrodes are present one of which is shown in Panel (d) in schematic form. Each electrode complex consists of inter-digitated electrodes with a gap of 20 micron in between. (e) shows light microscopic image of micro-fabricated gold electrodes (dark area) on cover-slip with biofilm grown on top. Image obtained with lowest magnification of light microscope using 5X objective lenses, image is reconstructed from 3 overlapping micrographs of the electrode surface. The relatively lighter regions are the channels between electrodes containing no gold but with biofilm grown on the insulating substratum (glass). Panel (f) is a schematic of the same measurement set-up in cross-section.(g) Photograph of transformation of initially pristine graphene during biofilm growth

To track the changes in electrical properties of the biofilm in presence and absence of graphene we have explored frequency dependent capacitance change. Capacitance (C) of biofilms is related to its charge storing ability. Many biological components contribute to charge storage e.g. proteins, polar head groups of lipids, chromophores, pigments etc. How interaction with graphene changes capacitance wouldl reveal the nature of possible interactions the biofilm may have with the graphene. Similarly, changes in impedance (G)^8–11^ are measured by applying sinusoidally oscillating voltage. G and C values are obtained as a function of frequency of oscillation.

In addition to this we also look for emergence of semiconducting properties. In pure graphene such properties are not expected^12^. Typically, semi-conducting materials possess an energy barrier or band gap (*V*_*B*_) between their valence electrons (closely bound) and conduction band (consisting of free electrons). Valence band electrons must be supplied with *energy* > *V*_*B*_ for transition to conduction band. A tight contact may be obtained between a metal (M) and semi-conductor(S), such that charge may flow from one to the other. The junction known as Schottky diode will have a detectable signature, namely a threshold voltage (*V*_*B*_).^13,14^ Current flow occurs only for voltages *V* > *V*_*B*_, which may be regarded as a threshold. Notably such property also arieses in the present case.

Graphene oxide (GO) is the oxidized form of graphene. Reduced graphene oxide (RGO) is obtained upon reduction of GO. Reduction removes oxygen from the material but dangling bonds remain open. Both RGO and GO have lesser conductivity than unmodified graphene. In situ reduced graphene oxide (RGO) incorporation has been used to modify microbial mat on MFC electrodes and increase its power output as well as in bio-photovoltaic devices.^15–19^

**Figure 2:**
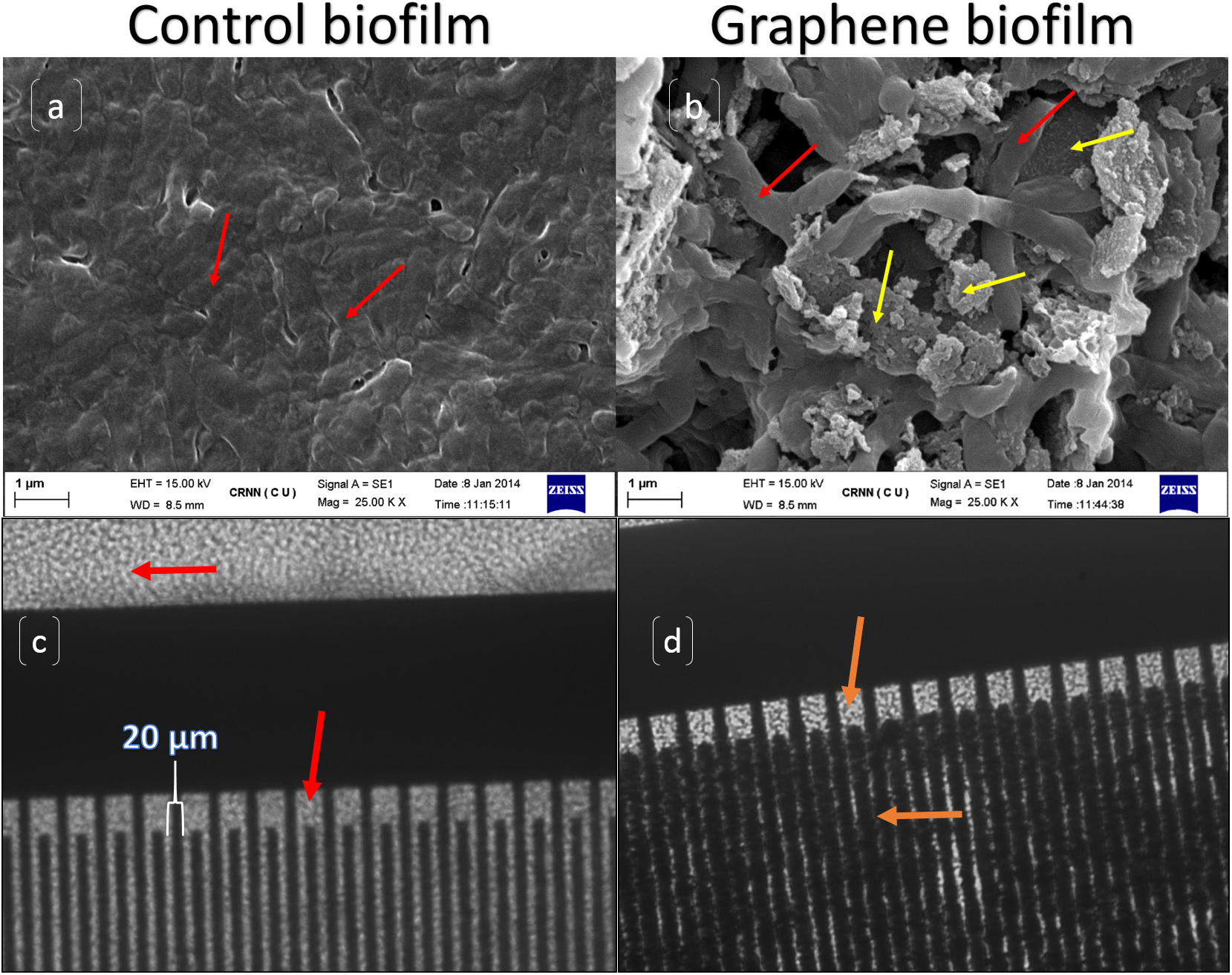
SEM images of (a) control biofilm compared to (b) graphene (yellow arrows) trapped by biofilm cells (red arrows) (c) Bright field microscopic images at 20X shows an IDE with control biofilm, i.e. without graphene. (finger width is 20 *µm*)(d) represents the fully grown and and partially peeled biofilm with incorporated graphene. Red arrows mark biofilm cells, orange arrows mark graphene incorporated composite.

## Results and discussion

The schematic diagram illustrating the graphene immobilization by biofilm is shown in the figure 1. It is difficult and costly to immobilize powdered forms of carbon nanomaterials (CNM) on device or electrode surface. Our work shows that biofilm may be used to form electrical contact between hydrophobic graphene and micro-fabricated electrodes as depicted in the figure 1. A functional electrical contact allows electron exchange between two different metals or between a metal and a semi-conductor.Studies generally do not vary amplitude of the AC voltage (RMS) has never been varied. By varying RMS voltage, points of convergence were obtained in the capacitance versus frequency spectra. We have studied how they differ in control and graphene incorporated biofilms. We study the electrical properties from 2 perspectives. We report biofilm induced band gap engineering of graphene. Secondly, use of an isosbestic point to monitor change in dielectric properties of biofilm by graphene.

### Electron microscopy

SEM images of graphene containing bio-composites show wrinkled graphene. Such few layered) wrinkled graphene (FIWG) is marked by yellow arrow (see figure 2 (a,b)) and is trapped by biofilm (indicated by the red arrow in the figure panels). No trapped sheet-like structures may be observed in electron micro-graphs of control samples. The average diameter of as incorporated platelets was between 1-2 *µm*, with a wide size distribution. Some larger wrinkled sheet like structures were observed, trapped underneath cells. It is to be noted that these are 11th day samples, by which time cell sloughing has occurred to a great extent thus decreasing the surface area covered by cells. Bacteria can be distinguished by their smoothness compared to rough surface of wrinkled graphene clusters.

#### Immobilization mechanism

From the SEM images, the immobilization appears to be due to trapping of FlWG underneath biofilms and in between layers of biofilm cells. Purple non-sulphur biofilms adhere strongly to a large variety of surfaces^20^ due to the presence of extra-cellular polysaccharides (EPS) and associated proteins. Thisis in consistence with previous studies on incorporation of graphene into bacterial membranes, by hydrophobic interaction with phospholipid tails.^21,22^ It may be noted,the release of FlWG from clumps, as initially added, is probably due to surfactants released during biofilm formation.^23^ Surfactants have been previously useful for obtaining single or few layered structures from multi-layered 2D material.^24^

### Raman characterization

Raman spectrum was obtained for the graphene that was used for this study. Raman spectrum (figure 3) obtained with red laser (632 nm) showed I_D_: I_G_ ratio of 1.259 and using the relation of this ratio to average distance between defects (*L*_*D*_) as in Cancado et al,^25^ *L*_*D*_ was determined to be 15.194 nm, which is greater than the average *L*_*D*_ for RGO which is around 3-4 nm.^26,27^ Hence our sample had relatively less defects compared to RGO and hence of higher hydrophobicity. Hydrophobicity is the important parameter that prevents interaction of graphene^28^ with most aqueous based biosystems.

The anomalous hump like 2D peak has been reported for few layer wrinkled graphene^29^ (FlWG), hence we may infer a part of the used graphene to contain FlWG. This is supported by the abnormally high FWHM of our samples of 65 cm^-1^ (G band) and 52 cm^-1^ (D band), as previously reported. In addition, the high intensity (almost same as *I*_*G*_) around 2D region suggests decoration of wrinkled forms on the surface of larger sheets (since FlWG does not have as high 2D) and is supported by the SEM images of trapped sheets (supplementary figure S2). Raman spectra of used graphene is important since FlWG are known to possess an intrinsic band gap in many cases.

The wrinkled nature is supported by another Raman spectrum obtained with 488 nm laser, which shows an average distance between defect of 10 nm (see supplementary figure S3) yet distance between edges of 4.358 nm (according to Tunistra-Koenig relation). Wrinkling and folding may be responsible for reducing the size by bringing the edges much closer to each other and increasing the effective defect density. Please note that due to folding, sheet like structure is modified into platelets or particulate matter.

**Figure 3:**
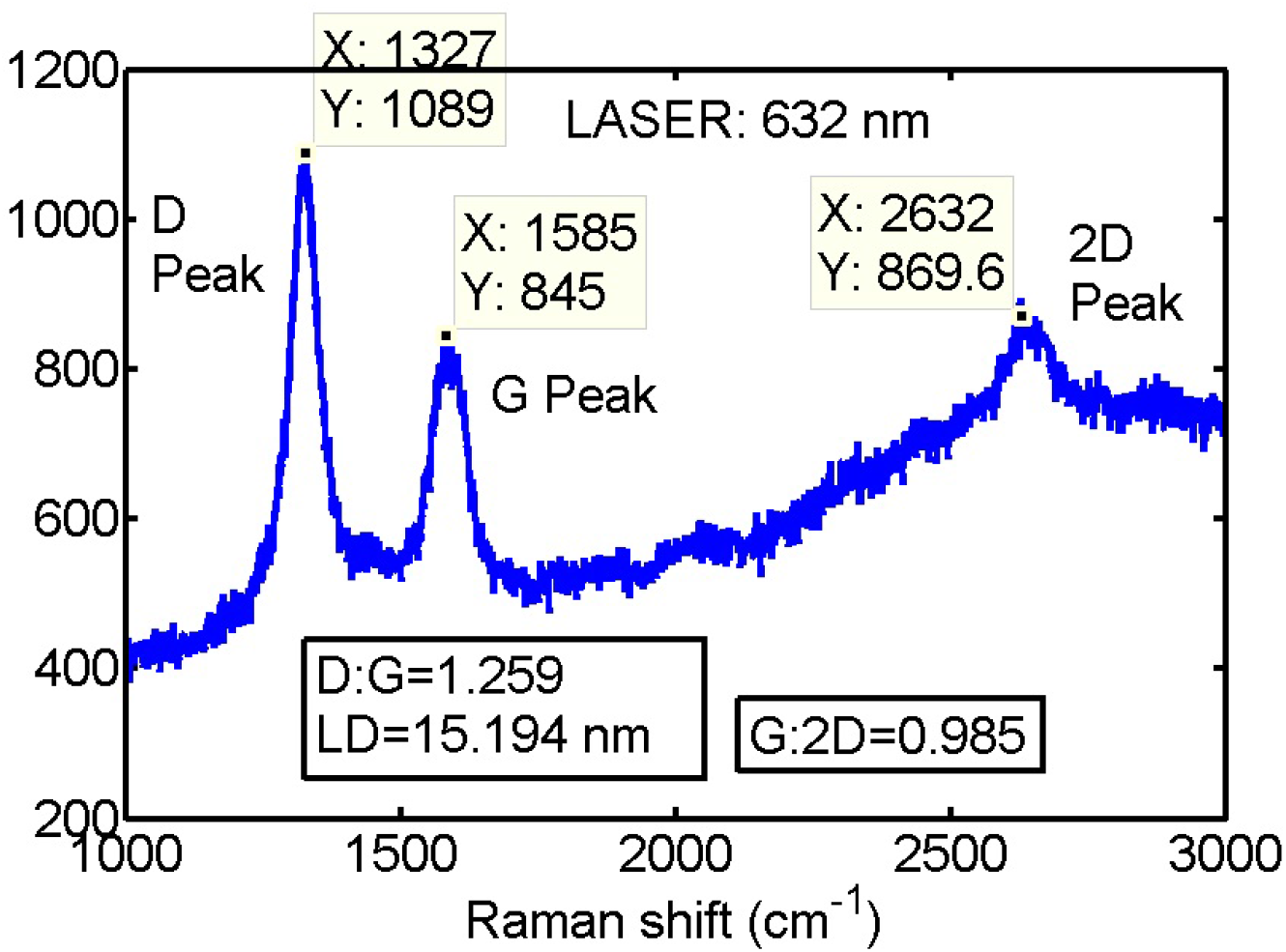
Raman spectrum of the graphene used in these experiments, obtained with red laser (632 nm). *I*_*D*_: *I*_*G*_ ratio was 1.259, and corresponds to *L*_*D*_ = 15.194 nm. 2D peak obtained showed a hump. FWHM was much greater than is typical for monolayered graphene. These features have been previously described for few layered wrinkled graphene.

Raman spectra of biofilm incorporated graphene could not be obtained since Raman signal from carbohydrates (see supplementary figure S4) masked the Raman signature of graphene. This is proof that graphene was effectively trapped within extra-cellular polysaccharide, so that its surface is not accessible to light.

### Biofilm formation assays

Addition of graphene has been shown to inhibit growth of bacteria in a number of studies.^21,30^ While growing the samples used earlier, we observed intenser pigmentation for graphene containing samples in our case.

Firstly, crystal violet staining of biofilms^31^ was performed as described in Materials and Methods and amount of biofilm biomass per falcon tube was measured. Biomass increased with the addition of graphene in growth media, compared to control. This growth promoting effect increased with increase in graphene dose. However for the highest dose (5.5 mg in 50 ml) biomass was lowered (see figure ‘4(a)). This probably indicates some sloughing off of films for the highest dose of graphene. The photograph in the inset shows the empty falcon tubes with biofilm grown on its walls. The numbers in brackets indicate quantity of graphene added to each falcon tube.

**Figure 4:**
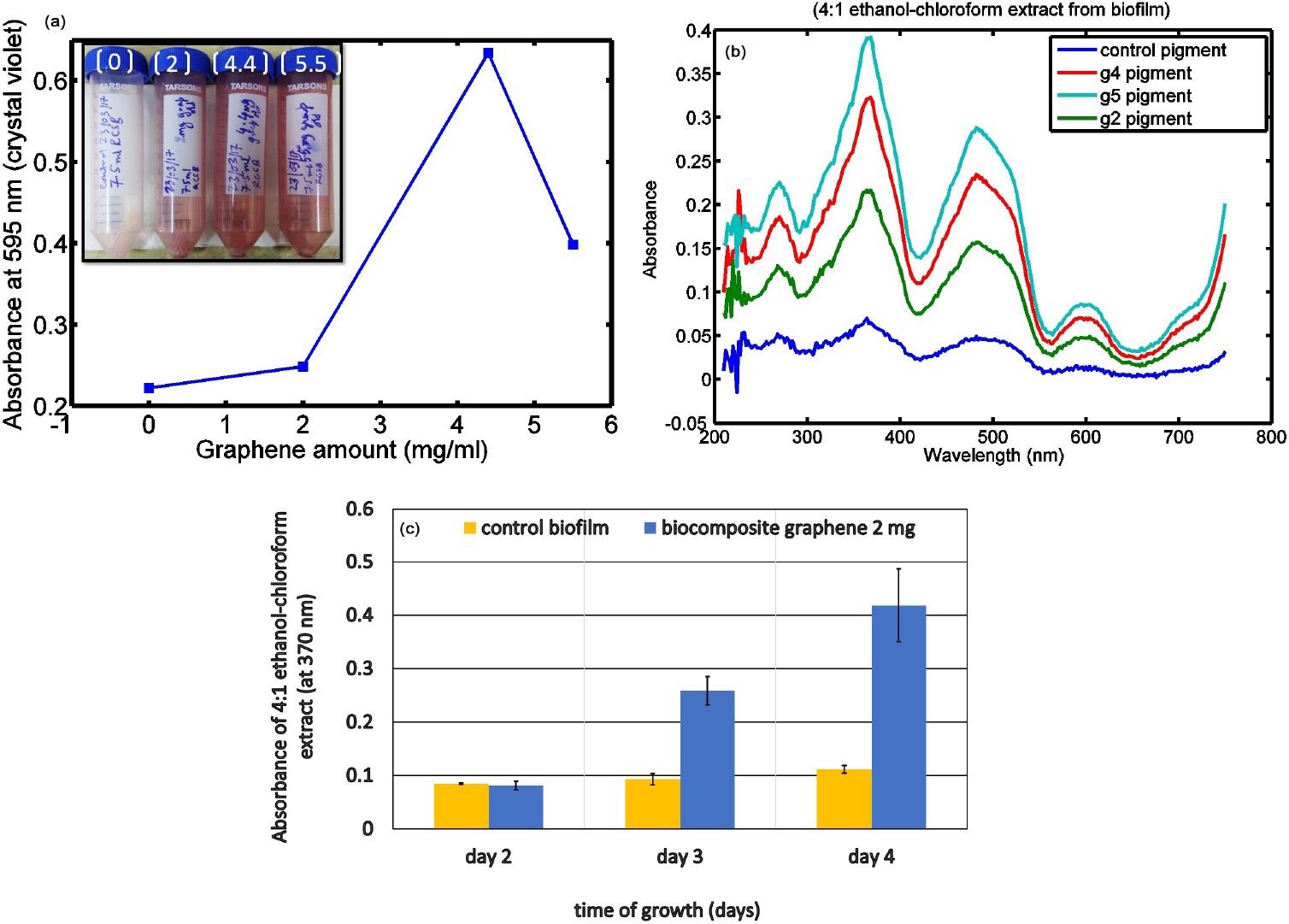
(a) Variation in biofilm biomass with increasing graphene dose, estimated with crystal violet staining assay. Inset shows photograph of biofilms adhered to falcon tube wall for each dose. (b) Variation in total pigment content of adhered biofilm with increasing graphene dose determined (c) Change in absorbance at 370 nm of organic extract of graphene-biofilm composite (2 mg graphene in 15 ml media) compared to Rhodobacter biofilm over 2-4 days of biofilm growth. Estimation of pigment is given by measuring the absorption spectra for organic extract (using chloroform: ethanol= 1: 4) from biofilms.

With respect to control, the total pigment content of biofilm on falcon tube walls increased with increasing graphene dose (see figure 4 (b)), for 4th day biofilms. This confirms increase of pigment content of biofilms when grown with graphene. Absorbance spectra of planktonic suspensions showed increasing O.D. with increasing graphene dose. This further confirms the growth promoting effects of graphene as well as indicating that there may be sloughing off from biofilm for the highest dose (5.5 mg).

The apparent growth promoting effect of graphene on RCSB biofilms was investigated by studying growth kinetics in presence and absense of graphene. From day 2 to day 4 of biofilm growth, absorbance at 370 nm showed a steeper increase (figure 4 (c)) when graphene was added (2 mg in 15 ml), compared to control. This proves that graphene promotes Rhodobacter biofilm growth.

### Effect of carbon source withdrawal

*Rhodobacter* was grown in a semi-synthetic medium where malic acid (4 mg/ml) was the major source of organic i.e. fixed carbon. To test whether graphene acts as alternative carbon source for *Rhodobacter*, biofilms were grown in various lesser (< 4 mg/ml) concentrations as well as in absence of malic acid (i.e. 0 mg/ml). Growth was allowed either in absence or presence of graphene. We have performed this experiment in order to probe the effect of a metabolite. The question is whether the presence of the metabolite would attenuate the graphene induced enhancement of bacterial growth. This is expected only if the bacteria use graphene as a carbon source. On the other hand if there is a co-opertaive effect beween the bacterial electron transport and the Dirac electron cloud. If this is the case we may expect a different result in which teh metabolism with synergestically increase in presence of graphene.

The results involving malic acid concentration variation ruled out the possibility of utilization of graphene as carbon source. It further demonstrated that overall pigment concentration increased in presence of graphene, but only when there was sufficient organic carbon source in the medium (see figure 5 a). The synergistic growth and pigment enhancing effects of graphene were further demonstrated by absorbance of the planktonic suspension at 392 nm (see figure 5 b). Both 370 nm (organic extract) and 392 nm (whole culture) absorbance peaks correspond to a fluorophore that has been reported in Rhodobacter.^32^ The fluorophore has excitation maximum at 395 nm and emits at 613 nm (see figure 5c).^32^ The fluorescence was additionally investigated. While graphene in all cases partially quenched the fluorescence, the increase in fluorescence with increasing malic acid concentration had a steeper slope when graphene was present. Hence the role of graphene has been to alter the photochemistry in the membrane of *Rhodobacter*. This points towards interaction between Dirac Fermions on the surface of graphene and the pigments associated with light energy harvesting in the purple non-sulphur bacteria. All the consequent emergent properties can be linked to this superposition of electron clouds.

**Figure 5:**
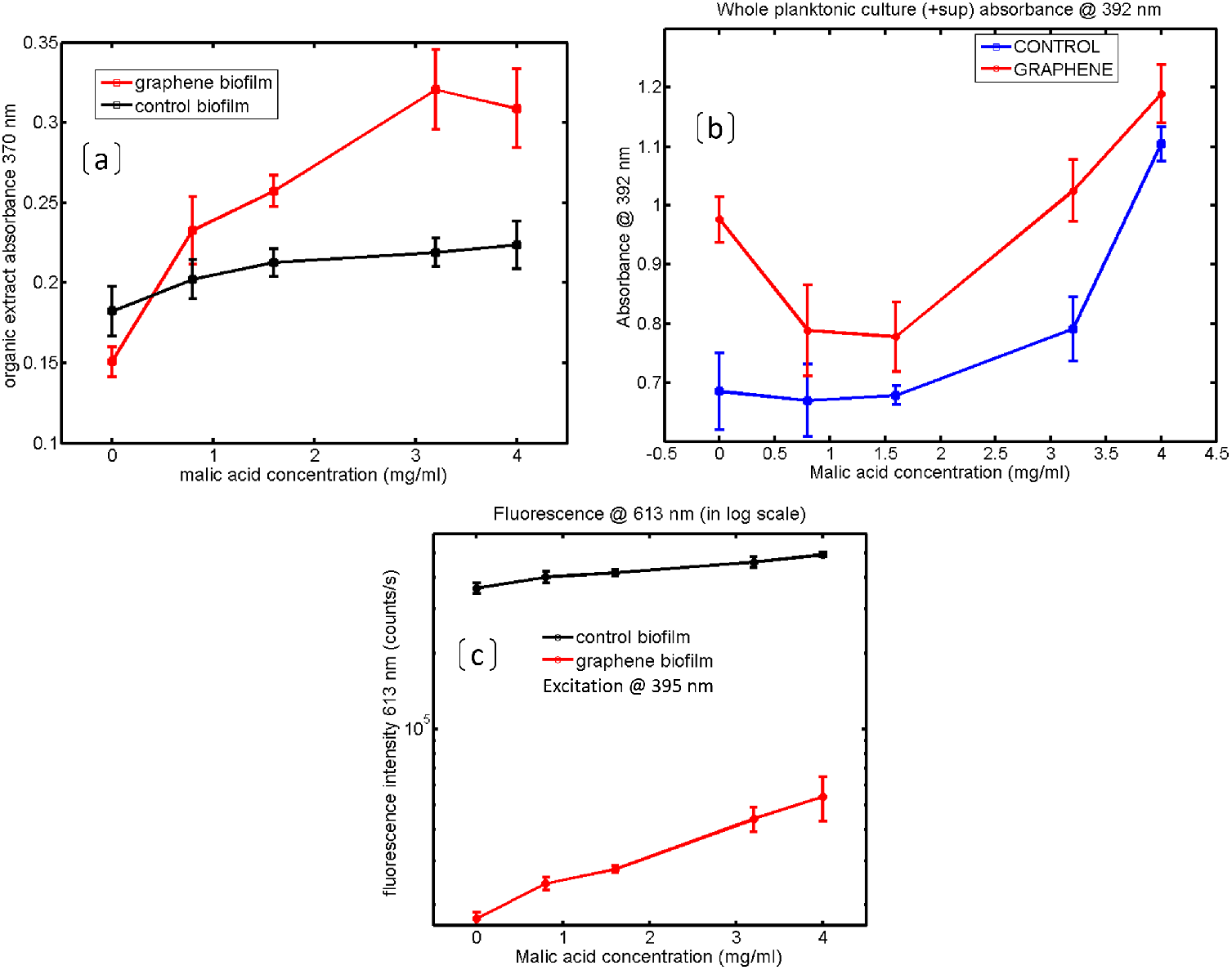
Variation in growth parameters for increasing concentrations of malic acid in growth medium, in presence and absense of graphene. (a) Absorbance of organic extract at 370 nm (b) Absorbance of planktonic suspension at 392 nm (c) Fluorescence at 613 nm with excitation at 395 nm.

### Electrical characterization

Our work shows the interaction between conducting graphene with charge carrier components in photosynthetic biofilms. IDE2 was used for current voltage measurements. IDE1 was used for capacitance measurements.

#### IV curves

Graphene incorporation into biofilm resulted in increased current flow, of the order of 10^−3^ A, for graphene dose of 4.4 mg in 50 ml (g4, see supplementary figure S6), compared to control, which gave current of the order of 10^−^11 A (see supplementary figure S6). For graphene dose of 2 mg in 50 ml (g2, see supplementary figure S6), current was of the order of 10^−6^ A, thus demonstrating that current through the biofilm depended on the quantity of graphene added to growth media. This proves graphene incorporation into biofilm. The reason behind plotting in log scale is that when samples differ in current conduction by a very large value, it is not possible to compare them in one figure.

Graphene incorporated biofilms exhibited sudden increase (of the order *X*10^2^ times) in current above a certain critical voltage (see 6 (a-c)). No such threshold was present in control biofilm. This voltage may be termed as a threshold. Presence of threshold voltage (*V*_*B*_) in IV curve indicates formation of a Schottky diode like junction (see Introduction for general description of Schottky diode^14^ between biocomposite and metal electrodes. The threshold voltage means that there is a conduction gate. Below *V*_*B*_, the gate is closed and charge flow is not allowed. Above *V*_*B*_, the conduction gate opens and current flow shows sudden increase.

Since graphene is the major conductive element in the biocomposite, we may conclude emergence of band gap in biofilm immobilized graphene.^3^

For g4 (see 6 (c)), threshold was symmetric (STH) for both positive and negative voltages (formally known as forward and reverse bias). In both cases, threshold occured for *V*_*B*_ > ±3.2 V.

However typical threshold was observed only when voltage was decreased from higher to lower values (i.e. voltage step-down followed the inequality *d*|V |/*dt* < 0). On the contrary when voltage was increased from 0 to higher positive or negative values, (i.e. if we follow the inequality *d*|*V* |/*dt* > 0) a hump in current conduction was observed (see Figure 6b,c,e and f). Hump was located near the Schottky threshold. Associated hysteresis in the IV curve (area within the curve) may be noted.

The gradient of current with respect to voltage (dI/dV, see Figure 6 (f)) i.e. conductance was plotted against corresponding voltage values. Conductance (G) versus voltage curve showed dependence of G on DC voltage (V). The conductunce-voltage curve showed prominent peaks at ±3.3*V*. Smaller peaks were observed at 0.4 and 1.6 V. The hump in current voltage curve correspond to the anomalous conductance peaks at ±3.3*V*. Voltage dependent conductance peaks^33^ are important since they may be an indication of ballistic transport.^34^ Note that, in this case, conductance peaks are occurring at room temperature (*T* >> 77*K*)^35^ and is linked to biofilm mediated graphene immobilization.

When measurements were performed over smaller voltage range -1 to 1 V (g4s, see supplementary figure S6 and S7), for g4, no threshold voltage was demonstrated (see figure 6(c)). Hysteretic properties (see supplementary figure S6 and S7) were scaled down, compared to -5 to 5 V.

**Figure 6:**
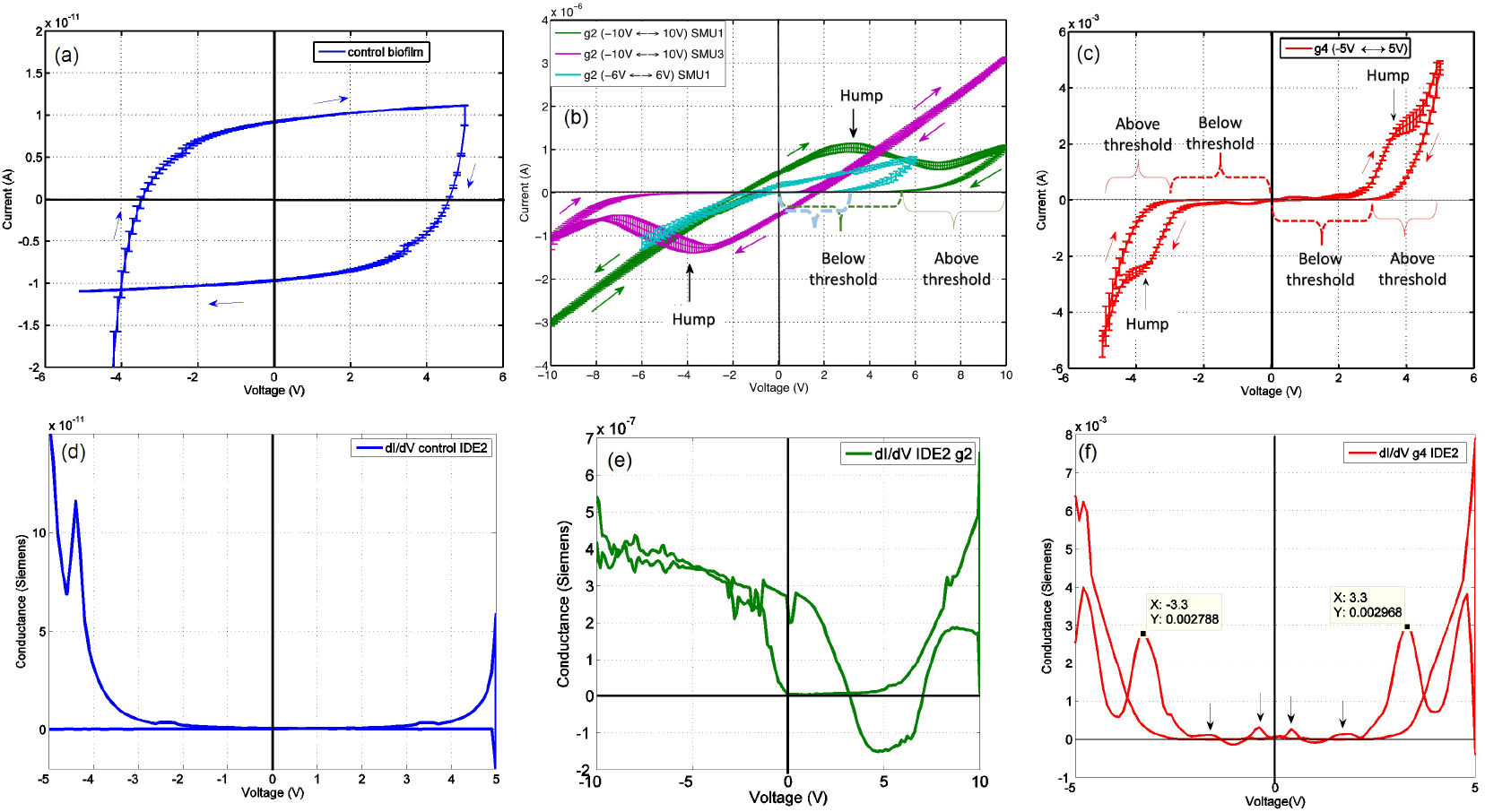
(a) IV curve for control biofilm showing hysteretic properties (area within the curve) of the biomaterial. (b) IV curve for biofilm-graphene composite grown with 2 mg graphene in 50 ml media (g2). Green and purple curves are for same sample but SMUs were interchanged. (c) IV curve for biofilm-graphene composite grown with 2 mg graphene in 50 ml media (g4). For (a-c) arrows indicate up or down sweep of voltage in each case. (d-f) Voltage dependent conductance derived from (dI/dV) at a given V for control biofilm, g2 and g4, respectively. The arrows in (f) show smaller conductance peaks along with the prominent peak at |*V*| = 3.3*V*, dI/dV were for obtained using mean current values.

For g2, threshold was asymmetric (ATH, see Figure 6 (b)) and present only when voltage was positive. Measurement with negative showed ohmic conduction (for green line). Thus one way Schottky diode was formed. This asymmetry may be linked to the asymmetry in current flow through a metal-semiconductor junction (see Introduction). This was confirmed by inverting the source measure units (SMU1 and SMU3) (see Figure 6 (b), purple line), which gave an inverted IV curve. This asymmetry was obtained only when low graphene dose was used (compare with Figure 6(c)).

For g2, similar to g4, only voltage step-down from higher values to 0 V (*d*|*V*|/*dt* < 0) resulted in conductive gating, i.e. dI/dV = 0. For voltage step-up (*d*|*V*|/*dt* > 0), capacitive gating was exhibited over a voltage range of 3.3 V to 7 V. By capacitive gating we refer to decrease in current with increase in voltage (dI/dV < 0, see Figure 7, for 3.3-7 V).

Measurement with smaller voltage range for g2 (g2s) resulted in shifted Schottky threshold along with disappearance of capacitive gating (blue line see Figure 6 (b)).

The conductance (dI/dV) versus voltage curve (see Figure 6(e), 7) for g2 shows the capacitance gate to be associated with negative conductance. Negative dI/dV implies flow of current opposite to the applied voltage implying emergence of capacitive component (see supplementary Figure S8). Area under the curve for this negative conductance region would give magnitude of capacitive current (see shaded area of figure 7). In figure 7, two regions are marked as conductive and capacitive gating. The regions show gates beyond which current rises steeply.

The bioelectronic device can be developed into a self-powered sensor device since it produces molecular *H*_2_.^36^ Production of *H*_2_ was achieved by immersing the sensor complex in fresh growth media and exposing to light(data not shown). Little bubbles appeared on the areas covered by biofilm. The self-produced *H*_2_ can be used to power the device into which the Schottky diode needs to function.

**Figure 7:**
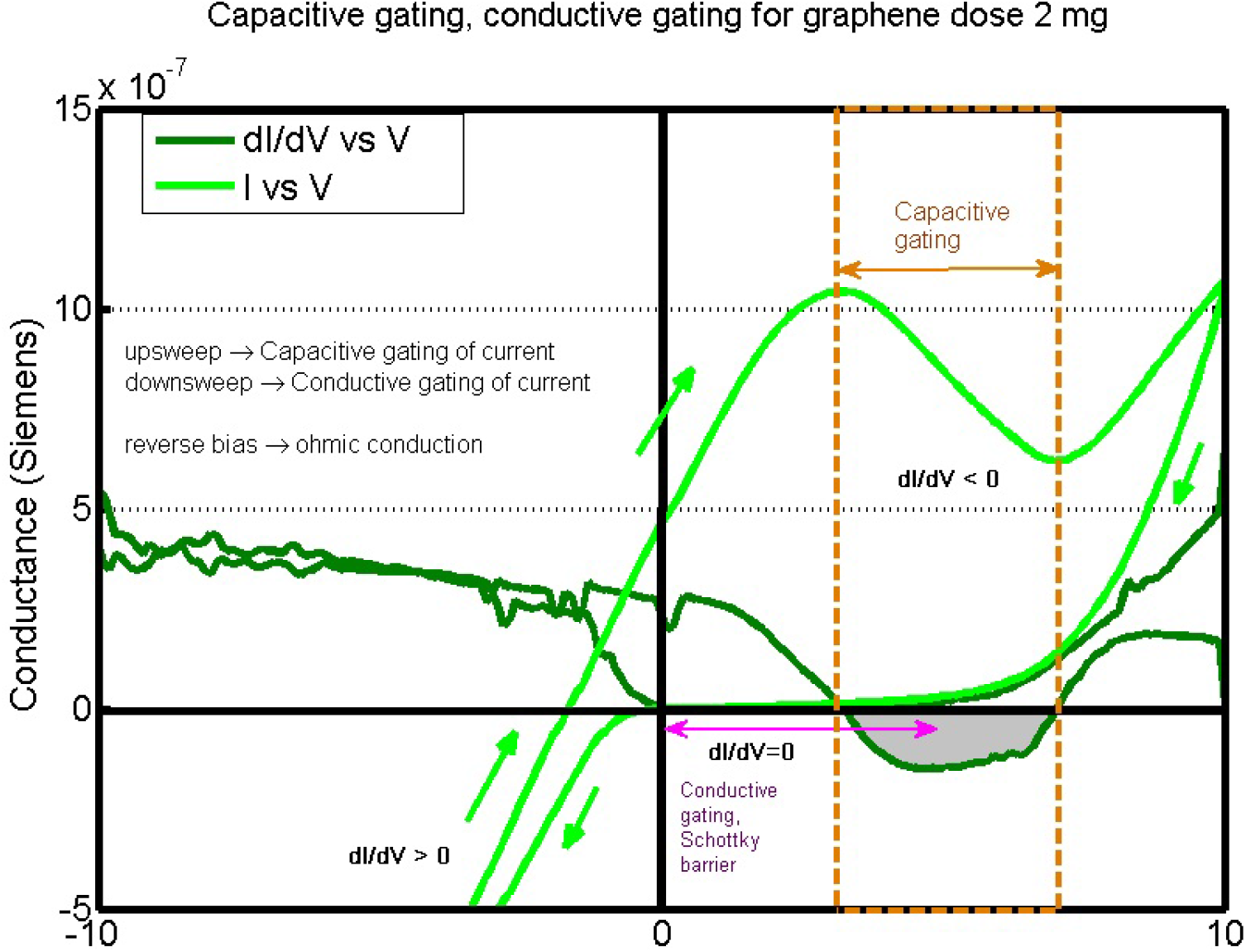
Current voltage curve for g2, overlayed with dI/dV versus voltage. Shows conductive gating and capacitive gating. Mean values of current were used.

#### Capacitance profiles

Capacitance profiles of both control (figure 8 (a)) and graphene incorporated biofilms (figure 8 (b)) exhibited co-dependence on frequency (*ν*) and RMS of AC voltage used for capacitance determination. Transition points were obtained such that capacitance values increased on one side (blue arrows) and decreased (red arrows) on other side of a particular frequency, upon RMS voltage variation. Spectra show an approximate convergent behavior near these transition points, i.e. values of capacitance come very close to each other and RMS dependence becomes negligible. We refer to these characteristic frequencies as capacitive isosbestics points, along the same line as spectrophotometric isosbestic points.^37^ Isosbestic points may be interpreted as signifying a two-state dielectric transition, A → B, in response to changing AC voltage amplitude. Isosbestic point is important in the present context because the media being studied are complex dielectrics, yet isosbestic points provide a specific benchmark for characterization and detection.

For control biofilms (figure 8 (a)), isosbestic points were located at 0.9 and 6 MHz. Increased RMS voltage resulted in increased capacitance at 0.9 MHz < *ν* < 6MHz (peak response at 5 MHz) with concomitant decrease for *ν* < 0.9*MHz* and *ν* > 6*MHz* (peak response at 80 KHz and 9 MHz). Graphene incorporated biofilms demonstrated changes in both isosbestic points, with the frequencies corresponding to isosbestic points exhibiting unusually high standard deviation of capacitance values for RMS=10 mV. The frequency at which peak response occured i.e. 5 MHz showed a clear shift to 4 MHz for higher graphene doses, see yellow circle in Figure 8 (b). It should be noted that with increasing RMS, bandpass properties of the dielectric changes and it transitions into an all-pass dielectric. We may translate the converging behavior into a frequency gate.

We explain the capacitive isosbestic point as occurring due to dipole alignment sensitivity to amplitude of sinusoidally varying voltage. Increasing RMS voltage resulted in increasing capacitance at < 6 MHz (peak response at 5 MHz) and decreasing capacitance at > 6 MHz (peak response at 9 MHz). This signifies that overall there are 2 dielectric states of the material (say P, Q) each of which show maximum alignment to the right and left of 6 MHz respectively. Increasing the RMS voltage converts P → Q. At 6 MHz, decrease in alignment of one state is exactly compensated for by increase in alignment of the other state. Depending upon whether it is a mole-to-mole P → Q conversion, unique isosbestic point is obtained, otherwise points of intersection shift.

A shifting convergence point signifies the generation of inter-mediates,^37^ during transition from one state to another. Addition of higher quantities of graphene may contribute towards generation of dielectric intermediates e.g. by adsorption of photosynthetic pigments. The high standard deviation of capacitance (at RMS=10 mV), selectively in the transition zone, strongly points towards role of graphene in affecting dipolar rearrangement in its vicinity.

Graphene incorporation during biofilm growth resulted in overall decrease in capacitance, compared with control biofilm. This indicates that there is formation of spanning clusters of conducting graphene between gold electrodes resulting in a discharge current, thus dissipating the charge stored. This is consistent with the increased current flow with DC voltage.

**Figure 8:**
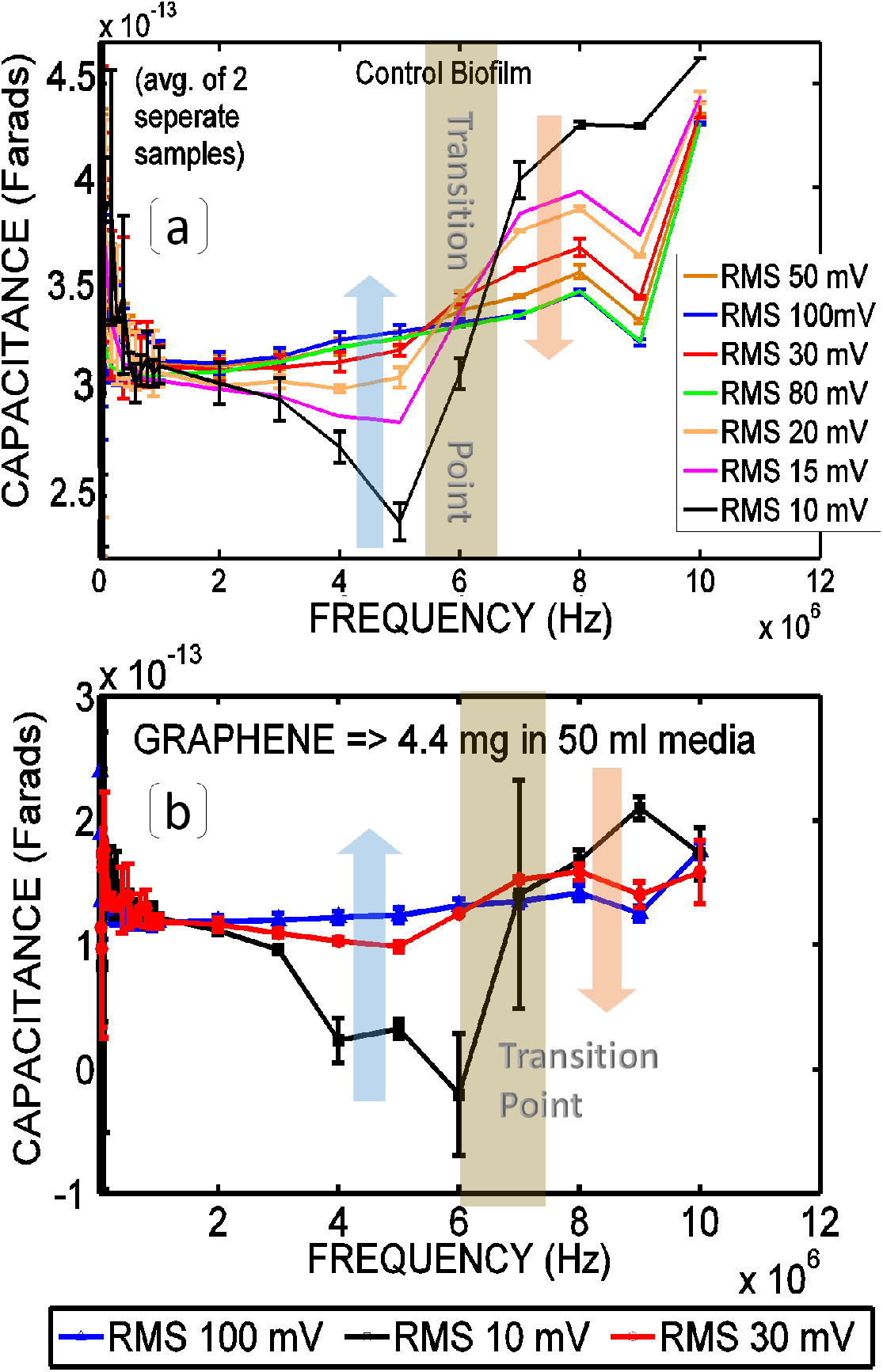
(a) Capacitance profiles of control biofilm. A transition point was obtained (marked by a grey zone in figure) such that capacitance increased with RMS (from 10 to 100 mV) on one side (blue arrow) and decreased with RMS on the other side (orange arrow). (b) Capacitance spectra of bio-composites (at RMS 10, 30 and 100 mV) for graphene (4.4 mg) added to 50 ml media.

The hysteretic property can be traced to biofilm. Hysteresis (area within the loop, see supplementary figure S6) increases when the amount of biofilm increases (see figure 4(a)).

#### Implication of the Schottky threshold

Graphene is a zero-band gap material meaning that there is no energy gap between its valence and conduction bands. Hence at room temperature bare graphene behaves as a metal. However, band gap is generated with strain induction in graphene or upon^12^ functionalization. Few layer wrinkled graphene is known to exhibit band gap.^29^

We explain the Schottky diode-like behavior as emerging due to (a) non-covalent functionalization of graphene by induction into lipid membranes of bacterial cells, due to hydrophobic interaction (b) graphene being held in a strained configuration due to trapping^38^ by extra-cellular polysaccharide (c) the intrinsic band gap of FlWG modified by the above two processes.

Presence of threshold points to band gap in entrapped graphene. Change in nature of threshold for different graphene doses points to role of graphene percolation through biofilm in determining behavior of entrapped graphene. This may be linked to the complex topology of pores in a biofilm.

All of the above processes induce curvature in otherwise flat graphene monolayer.^3^ Introduction of curvature is associated with generation of chiral vector. This curvature is analogous to the curvature in case of carbon nanotubes. Curvature in CNTs is described by its chiral vector. Chiral vectors affect band gap.^39,40^

Hence our next hypothesis from this work is that relative amounts of graphene and bacteria in a biofilm-graphene composite(compare figures 6 (b) and (c)) controls the nature of curvature (represented by chiral vectors) and hence the emergent band gap of trapped graphene. This would explain the difference in nature of Schottky barrier for the two doses of graphene. Thus biofilm growth modulation may be used to obtain tuneable metal-semiconductor junctions.

Opening of band gap provides opportunity for sensor design. Schottky diodes have been used as sensors.^41^ The present work shows biofilm controlled Schottky diode. Any receptor-ligand binding within biofilm, would cause shift in the junction threshold. Thus a new paradigm opens where a living material may be used as sensing platform.

## Conclusions

Interaction between graphene and purple non-sulphur bacteria is viewed from the perspective of interaction between two entirely different classes of electron clouds. For graphene, there is shift from conducting to semi-conducting property and for bacteria there is increase in is metabolism. In addition, there are some emergent properties present neither in biofilm nor in graphene. Schottky junction formation and its dependence on graphene biofilm stoichiometry is an example for such emergent property. There is an indication that the approach can be exploited for tuning band gap, which may lead to fabrication of flexible, semi-conducting biomaterial.

One notable observation was presence of electrical isosbestic point. In spectroscopy, isosbestic points denote two-state transitions. In the present case of graphene biofilm, the implication of existence of isosbestic point may be intriguing. Essentially it means there is a single set of frequencies that corresponds to di-electric transitions specific for the bio-conducting material. This may be a powerful tool for biofilm classification.

There are is one important questions that may be taken up in future: how two dimensional semi-conducting materials, with intrinsic band gap, (e.g. MoS2 and boron nitride) may exhibit modified band gap when entrapped within biofilm.

## Acknowledgements

This work was supported by DBT (BT/PR3957/NNT/28/659/2013).

## References

(1) Wehling, T.; Black-Schaffer, A. M.; Balatsky, A. V. Advances in Physics 2014, 63, 1–76.

(2) Dvorak, M.; Oswald, W.; Wu, Z. Scientific reports 2013, 3, 2289.

(3) Deng, S.; Gao, E.; Wang, Y.; Sen, S.; Sreenivasan, S. T.; Behura, S.; Král, P.; Xu, Z.; Berry, V. ACS nano 2016, 10, 8403–8412.

(4) Wilking, J. N.; Angelini, T. E.; Seminara, A.; Brenner, M. P.; Weitz, D. A. MRS bulletin 2011, 36, 385–391.

(5) Diner, B. A.; Petrouleas, V.; Wendoloski, J. J. Physiologia Plantarum 1991, 81, 423–436.

(6) Rezanka, T.; Siristova, L.; Melzoch, K.; Sigler, K. Mini-Reviews in Organic Chemistry 2010, 7, 300–313.

(7) Reguera, G.; Nevin, K. P.; Nicoll, J. S.; Covalla, S. F.; Woodard, T. L.; Lovley, D. R. Applied and environmental microbiology 2006, 72, 7345–7348.

(8) Malvankar, N. S.; Mester, T.; Tuominen, M. T.; Lovley, D. R. ChemPhysChem 2012, 13, 463–468.

(9) He, Z.; Mansfeld, F. Energy & Environmental Science 2009, 2, 215–219.

(10) Marsili, E.; Rollefson, J. B.; Baron, D. B.; Hozalski, R. M.; Bond, D. R. Applied and environmental microbiology 2008, 74, 7329–7337.

(11) Kim, T.; Kang, J.; Lee, J.-H.; Yoon, J. Water research 2011, 45, 4615–4622.

(12) Han, M. Y.; Ozyilmaz, B.; Zhang, Y.; Kim, P. Physical review letters 2007, 98, 206805.

(13) Nigro, M. A.; Faggio, G.; Fedi, F.; Polichetti, T.; Miglietta, M. L.; Massera, E.; Di Francia, G.; Ricciardella, F. Cross interference effects between water and NH 3 on a sensor based on graphene/silicon Schottky diode. In AISEM Annual Conference, 2015 XVIII; 2015.

(14) Svensson, J.; Campbell, E. E. B. Journal of Applied Physics 2011, 110, 111101.

(15) Ng, F.-L.; Jaafar, M. M.; Phang, S.-M.; Chan, Z.; Salleh, N. A.; Azmi, S. Z.; Yunus, K.; Fisher, A. C.; Periasamy, V. Scientific reports 2014, 4, 7562.

(16) Zhuang, L.; Yuan, Y.; Yang, G.; Zhou, S. Electrochemistry Communications 2012, 21, 69–72.

(17) Yuan, Y.; Zhou, S.; Zhao, B.; Zhuang, L.; Wang, Y. Bioresource technology 2012, 116, 453–458.

(18) Yong, Y.-C.; Yu, Y.-Y.; Zhang, X.; Song, H. Angewandte Chemie International Edition 2014, 53, 4480–4483.

(19) Liu, J.; Qiao, Y.; Guo, C. X.; Lim, S.; Song, H.; Li, C. M. Bioresource technology 2012, 114, 275–280.

(20) Garrett, T. R.; Bhakoo, M.; Zhang, Z. Progress in Natural Science 2008, 18, 1049–1056.

(21) Tu, Y. et al. Nature nanotechnology 2013, 8, 594–601.

(22) Frost, R.; Jonsson, G. E.; Chakarov, D.; Svedhem, S.; Kasemo, B. Nano letters 2012, 12, 3356–3362.

(23) Davey, M. E.; Caiazza, N. C.; O’Toole, G. A. Journal of bacteriology 2003, 185, 1027–1036.

(24) Trapalis, A.; Todorova, N.; Giannakopoulou, T.; Boukos, N.; Speliotis, T.; Dimotikali, D.; Yu, J. Applied Catalysis B: Environmental 2016, 180, 637–647.

(25) Cancado, L. G.; Jorio, A.; Ferreira, E. M.; Stavale, F.; Achete, C.; Capaz, R.; Moutinho, M.; Lombardo, A.; Kulmala, T.; Ferrari, A. Nano letters 2011, 11, 3190–3196.

(26) Eigler, S.; Dotzer, C.; Hirsch, A. Carbon 2012, 50, 3666–3673.

(27) Grimm, S. B.; Schweiger, M.; Eigler, S.; Zaumseil, J. The Journal of Physical Chemistry C 2016.

(28) Mao, S.; Lu, G.; Yu, K.; Bo, Z.; Chen, J. Advanced materials 2010, 22, 3521–3526.

(29) Kaniyoor, A.; Ramaprabhu, S. AIP Advances 2012, 2, 032183.

(30) Hu, W.; Peng, C.; Luo, W.; Lv, M.; Li, X.; Li, D.; Huang, Q.; Fan, C. ACS nano 2010, 4, 4317–4323.

(31) Merritt, J. H.; Kadouri, D. E.; O’Toole, G. A. Current protocols in microbiology 2005, 1B–1.

(32) Bose, A.; Raja, S. O.; Chowdhury, R.; Nandi, S.; Ray, S.; Bhattacharyya, K.; Dasgupta, A. K. bioRxiv 2017, 167122.

(33) Ilin, A. I.; Aparshina, L. I.; Dubonos, S. V.; Tolkunov, B. N. Russian Microelectronics 2001, 30, 17–21.

(34) Rickhaus, P.; Liu, M.-H.; Makk, P.; Maurand, R.; Hess, S.; Zihlmann, S.; Weiss, M.; Richter, K.; Schönenberger, C. Nano letters 2015.

(35) Poncharal, P.; Berger, C.; Yi, Y.; Wang, Z.; de Heer, W. A. arXiv preprint cond-mat/0211515 2002.

(36) Keskin, T.; Hallenbeck, P. C. 2012, 313–340.

(37) Berlett, B. S.; Levine, R. L.; Stadtman, E. R. Analytical biochemistry 2000, 287, 329–333.

(38) Kempaiah, R.; Salgado, S.; Chung, W. L.; Maheshwari, V. Chem. Commun. 2011, 47, 11480–11482.

(39) Timp, G. Springer, New York.

(40) Dresselhaus, M.; Dresselhaus, G.; Saito, R. Physical Review B 1992, 45, 6234.

(41) Kim, C.; Lee, J.; Lee, Y.; Cho, N.-I.; Kim, D. Sensors and Actuators B: Chemical 2000, 66, 116–118.

